# Time Machine: Can a Dye from 1928 be Re-Purposed for Modern, Fluorescence-Based Detection of Amyloid-Like Fibrils?

**DOI:** 10.1101/665364

**Authors:** O. I. Antimonova, N. A. Grudinina, V. V. Egorov, V. V. Ilyin, Y. A. Zabrodskaya, E. S. Ramsay, V. A. Polukeev, M. M. Shavlovsky

## Abstract

This manuscript describes the chemical synthesis of a compound similar to fluorene and Congo red, including characterization of its spectral properties. It was shown that the dye, during interaction with amyloid-like fibrils of several proteins (lysozyme, insulin, and beta-2-microglobulin), has the ability to change fluorescent spectrum. In contrast, monomeric forms of these proteins did not induce significant spectral changes.

## Introduction

To date, more than 30 human diseases associated with the formation of tissue amyloid deposits (amyloidoses) are known. In accordance with modern nomenclature, the type of amyloidosis is determined by the fibrillogenic protein involved in pathogenesis [1]. Since the introduction of terms such as “amyloid” and “amyloidosis” into the practice of medicine in the middle of the 19th century by Rudolf Virkhov, the molecular bases of these conditions have been well-studied, yet problems related to diagnosis and therapy still surround them. The “gold standard” for detection of amyloid deposits is the Congo red azo dye tissue sample assay, as introduced into practice by Bennhold H. in 1922. As noted in guidelines [1], the Congo red assay is the most reliable method of amyloid deposit detection, yet in some cases it is inapplicable. One of the drawbacks of the assay is the need to use green birefringence, rather than fluorescence for the visualization of amyloid deposits. The use of fluorescent dyes, such as thioflavins [2] and a new thiophene derivative [3], does not always allow unambiguous amyloid identification. Therefore, the search for new, more convenient, ways to detect amyloid in various tissues is an ongoing concern. In this context, a promising approach is to test new fluorescent dyes with structural similarities to Congo red. We have shown data on a diaminofluorene dye, Analogue of Fluorene (AF), including its efficiency of amyloid deposit detection in tissues [4]. It was not demonstrated, however, if it is universally suitable for amyloid fibril detection under *in vitro* conditions. This manuscript is dedicated to better understanding AF *in vitro* and to closing the gap between our histochemical and spectral knowledge. We present here analysis of AF’s synthesis and spectral properties in complexes, including absorption and fluorescence properties.

## Materials and methods

### Materials

All chemicals used in dye synthesis, including buffers and reagents, were purchased from Sigma (USA).

### Methods

#### ‘Analogue of Fluorene’ (AF) Synthesis

The Congo red homologue ‘Analogue of Fluorene’ (2,7-bis(1-amino-4-sulfonaphthyl-2-azo)fluorine) was synthesized as follows. 2,7-diaminofluorene dihydrochloride (269 mg; 1 mM final) was dissolved with stirring and ice-cooling in 12 ml of water:HCl (10:1) mixture. An aqueous NaNO_2_ solution (140 mg; 2 mM final) was added drop-wise in the presence of a catalytic amount of crystalline copper sulfate, and the mixture was stirred for 20 minutes (Solution A, bis-diazonium salt). Separately, a solution of disodium salt of naphthionic acid (2 mM final, Solution B) was prepared as follows. A suspension of naphthionic acid (446 mg; 2 mM final) was prepared in water. 10N NaOH was added drop-wise until complete dissolution of naphthionic acid. Solution A was added (with stirring and cooling) to 10 ml of Solution B. After that, tripartite sodium acetate (1 g; 7.35 mM final) was added, and the mixture was stirred for 30 minutes until the complete formation of the desired dark red azo coupling product. All reactions were carried out under nitrogen. Sodium chloride was added to the resulting product solution until saturation, and the precipitate was separated by centrifugation (30 min, 10000 g), followed by dissolution in a minimal volume of DMSO. The dye was precipitated with 96% ethanol and dissolved in a small amount of phosphate buffer (pH 8.0), followed by HPLC purification. According to mass spectrometry (data not shown), the molecular weight of AF corresponded to expectations (including identifiable peaks related to the disodium salt and the acid form of the dye).

#### Analytical and preparative HPLC

Reversed-phase column chromatography was used (HPLC YMC Triart C18, 250×20 mm, 5 μm, 12 nm). Chromatography was carried out in gradient mode. The mobile phase had the composition AB, wherein: A was a pure water; B was acetonitrile. For elution, mobile phase composition was as follows: 99% A + 1% B (200 ml); gradient transition from 99% A + 1% B to 40% A + 60% B (400 ml); and then 40% A + 60% B (100 ml). For the assay, an AF solution (in buffer A) at a concentration of 0.7 mg/ml was used. The detection was carried out using a UV detector at 214, 254, 360, 480, and 600 nm wavelengths. The peak corresponding to the target component had a retention time of 49.0 min (retention volume 410 ml). Integration of peak areas allowed us to conclude that the purity of our AF synthesis was approximately 95%.

#### Fibril preparation

Preparation of fibrils was performed in the same manner as described earlier for beta-2-microglobulin [5], lysozyme [5], and insulin [6]. All fibril solutions were tested using thioflavin T assay.

#### Thioflavin T assay

The method used here is a modification of an earlier assay [7]. Fibril suspensions or protein solutions (2.5 μM lysozyme; 0.17 μM beta-2-microglobulin; or 0.72 μM insulin) were added to a 25 μM solution of Thioflavin T in 50 mM phosphate buffer (pH 8.0). The fluorescence spectrum was recorded with a Hitachi F-7000 FL (Japan) spectrophotometer using 440 nm excitation (5 nm slit width) and fluorescence detection at 450-700 nm (5 nm slit width).

#### AF absorption spectrum

To a 10 μM solution of AF (in 50 mM phosphate buffer, pH 8.0), 5 μM of protein solution or fibril suspensions (to be tested) were added, and the absorption spectrum was recorded on a NanoDrop 2000 (Thermo, USA) spectrophotometer at 350-750 nm.

#### AF fluorescence assay

To a 2.5 μM solution of AF in buffer (in 50 mM phosphate buffer, pH 8.0), 2.5 μM of protein (protein solution or fibril suspension) was added, and the fluorescence spectrum was recorded on a Hitachi F-7000 FL (Japan) spectrophotometer using 530 nm excitation (5 nm slit width) and fluorescence detection at wavelengths of 570-700 nm (5 nm slit width).

## Results and discussion

The structure of AF (analogue of fluorene) is shown in Figure 1a. It was first described as a fabric dye [8], and it has never been used as a fluorescent probe for the determination of amyloid-like fibrils, despite its structural similarity with Congo red (Figure 1b). AF’s affinity for protein aggregates and ability to bind fibrils have been described [9,10]. The dye was synthesized using an azo coupling reaction with naphthionic acid disodium salt and the diazo derivative of diaminofluorene. In our previous work [5], we have shown that commercial Congo red reagents contain significant amounts of impurities. Here, we used our previously developed method for dye production. A decision was made to add an HPLC step, both as a final purification and as a convenient means to evaluate impurity levels.

**Figure 1.**
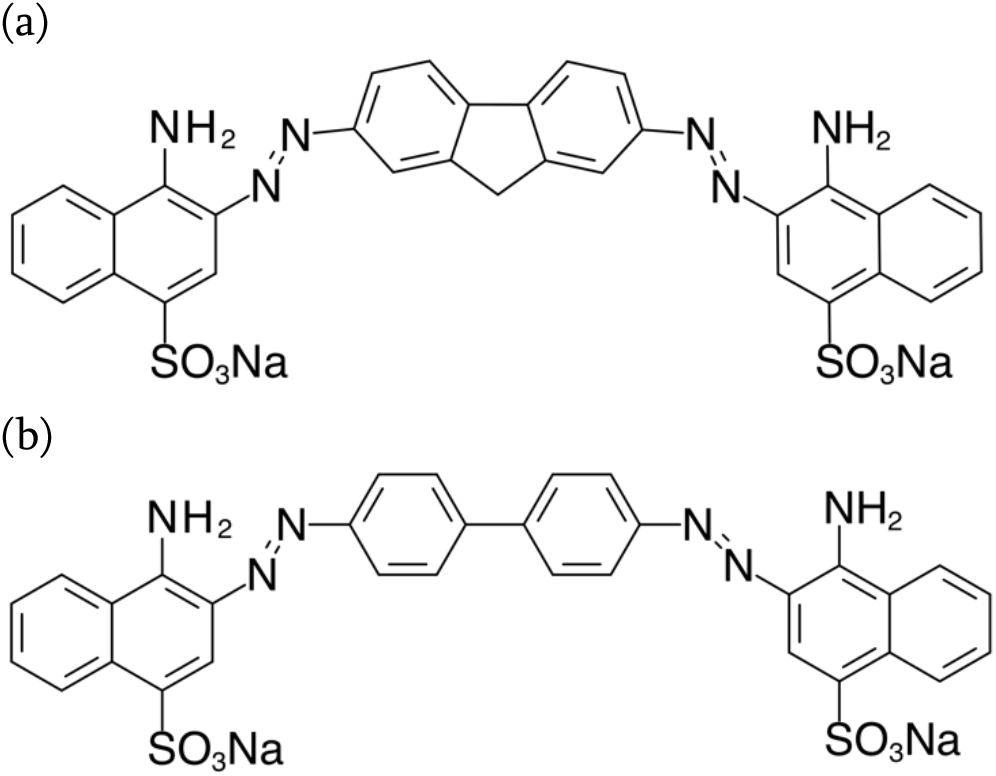
Chemical structure of (a) AF dye and (b) Congo Red.

Figure 2 shows the absorption spectrum of purified AF aqueous solutions at pH 2.0, 5.0, and 8.0. The spectrum is characterized by one maximum in the long-wavelength region, and the maximum extinction coefficient was observed at pH 8.0.

**Figure 2.**
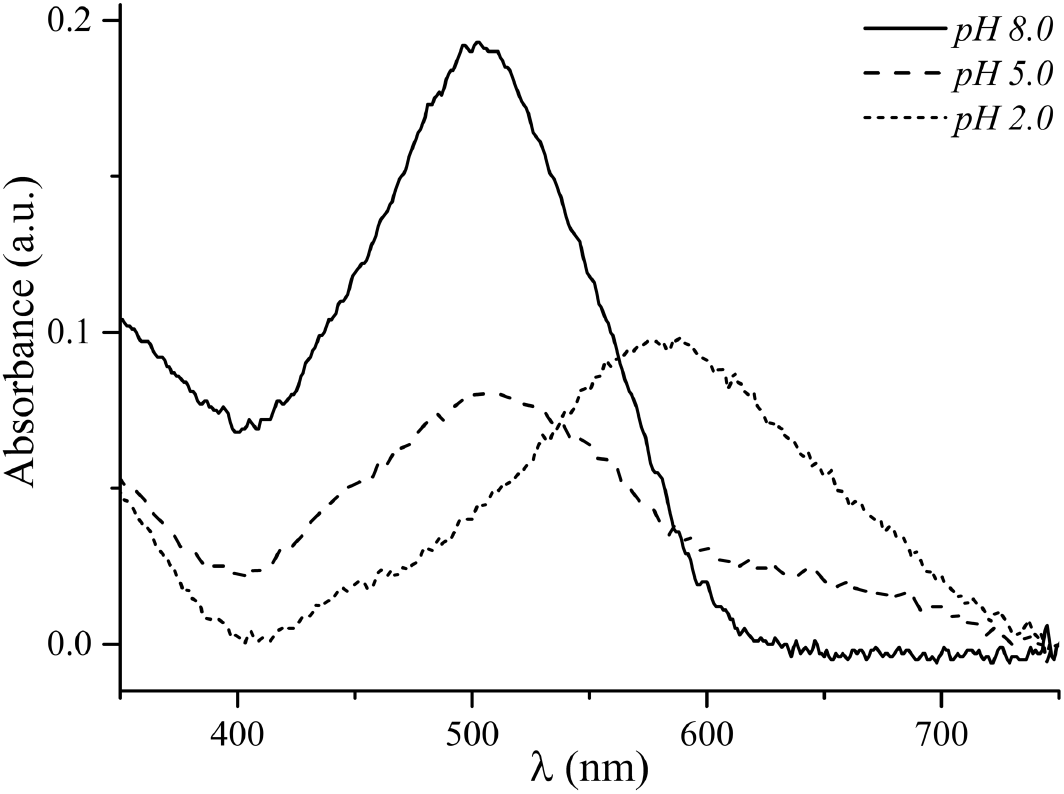
AF absorption spectra at different pH values.

The effect of soluble and fibrillar forms of fibrillogenesis-capable proteins on the AF absorption spectrum in solutions at pH 8.0 were studied. The presence of fibrils in study samples was verified using Thioflavin T (ThT) assay. Figure 3 shows the ThT emission spectra of the protein fibril and monomer solutions tested.

**Figure 3.**
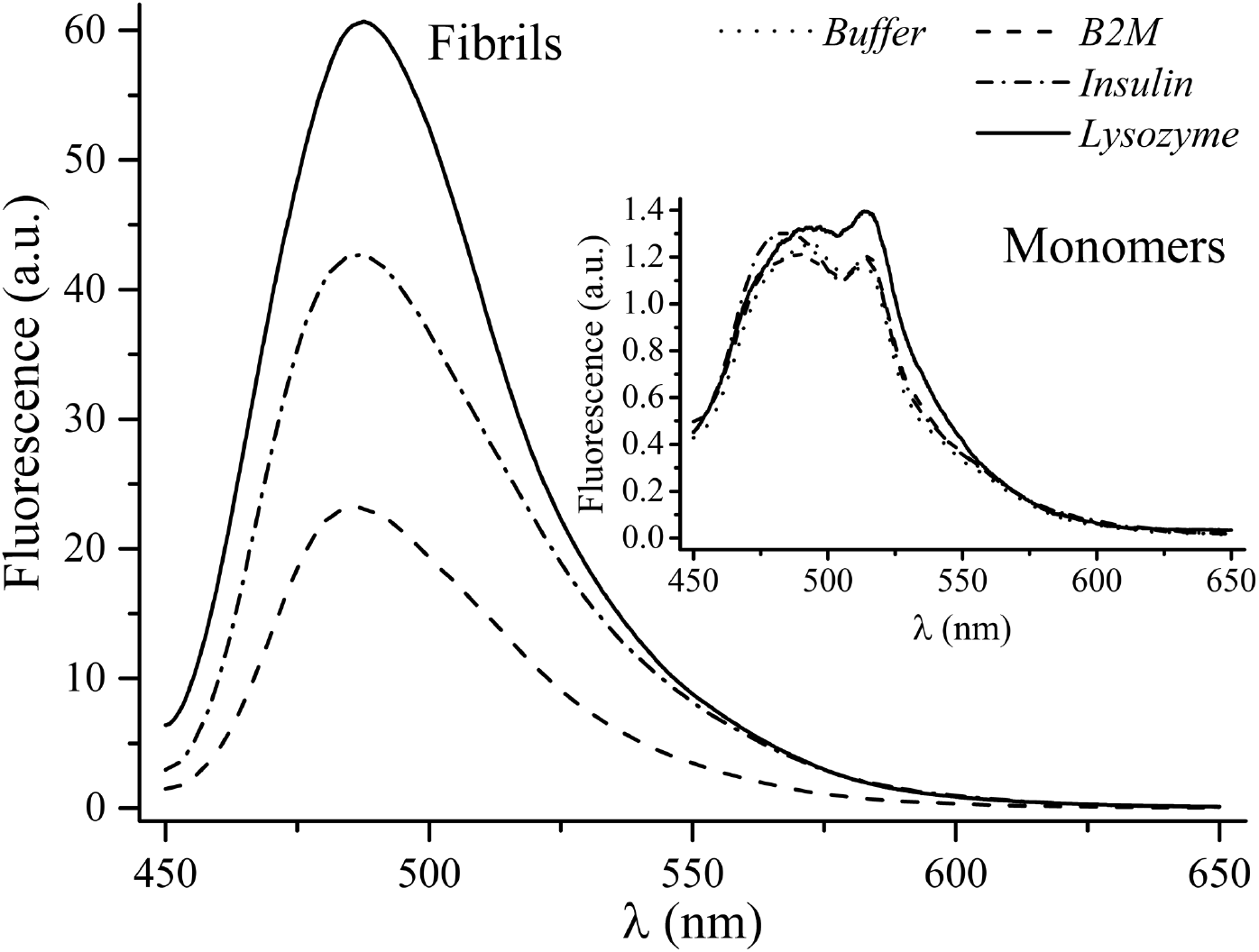
ThT complex fluorescence spectra of different soluble and fibrillar proteins.

ThT assay shows a characteristic increase in quantum yield in the presence of amyloid-like fibrils.

The tested protein solutions and fibril suspensions (Fig. 3) were the study materials used for further AF dye testing experiments. The addition of protein fibrils to AF dye solutions (at pH 8.0) revealed shifts in absorption maxima: from 500 nm to 514 nm for lysozyme fibrils; from 500 nm to 525 nm for insulin fibrils; and from 500 nm to 524 nm for beta-2 microglobulin fibrils. Increases in absorption intensity were also seen (Figure 4).

**Figure 4.**
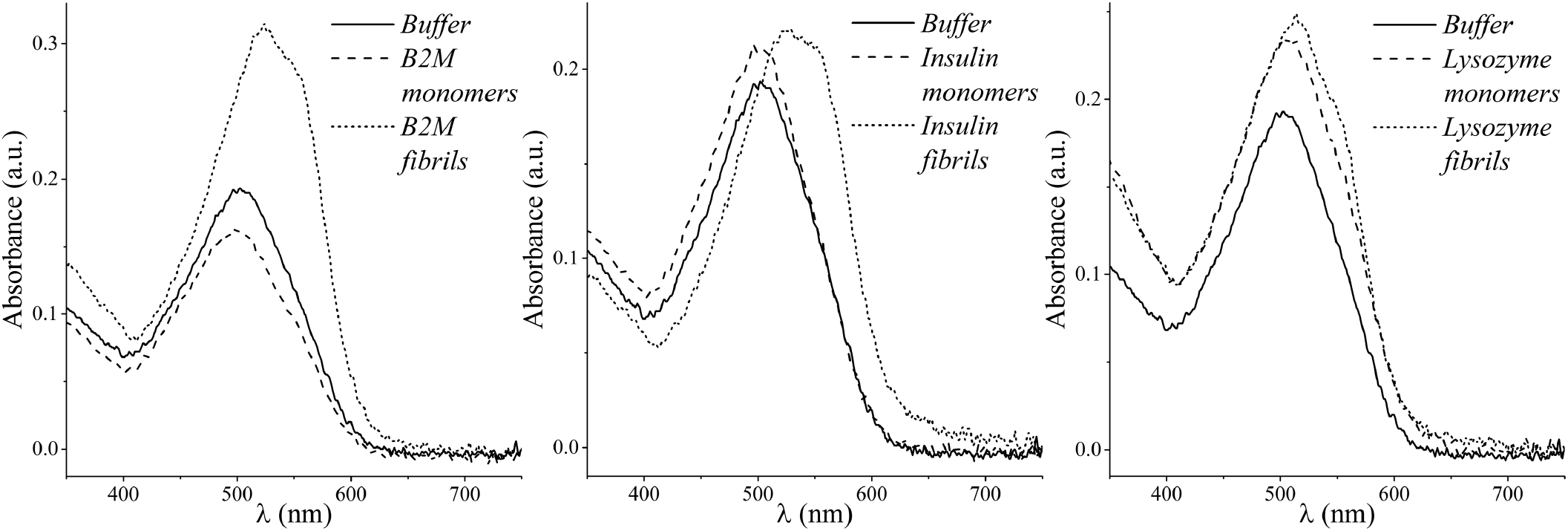
AF absorption spectra with different proteins in soluble form an as fibrils.

The shift of the absorption maximum to the right resembles the Congo red shift [5], and may be due to similar mechanisms seen in AF and CR binding. AF dye in solution shows red fluorescence. According to fluorimetry data of the AF-soluble proteins mixes, the excitation maximum corresponds to 530 nm; the emission maximums are 590 nm for non-fibrillar insulin and 650 nm for other proteins tested. As shown in Figure 5, AF-fibril suspensions featured the same excitation maxima (590 nm), and fluorescence intensities increased when dye/protein complexes were formed for all of the amyloidogenic proteins studied (insulin, beta-2 microglobulin, and lysozyme).

**Figure 5.**
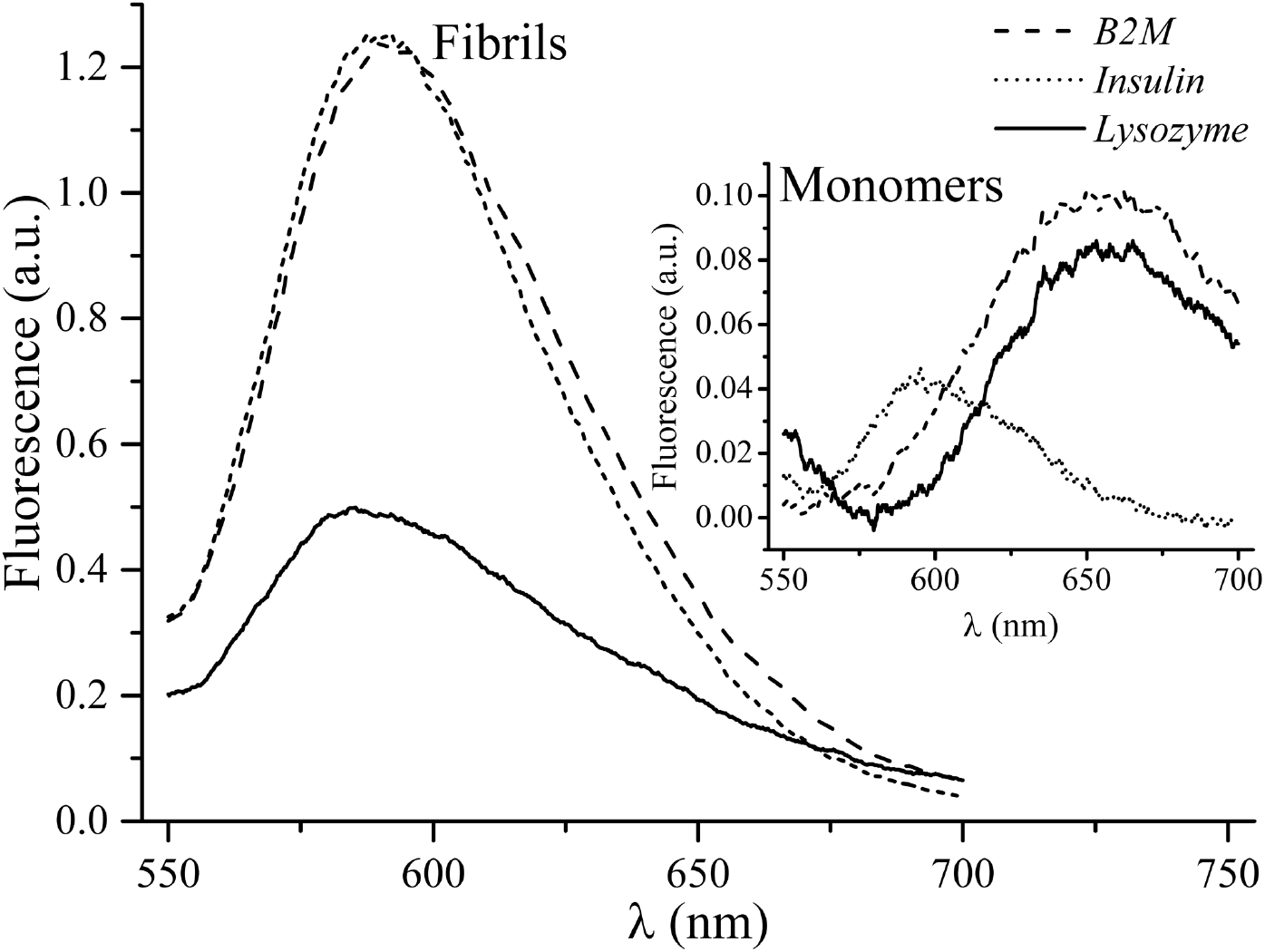
Fluorescence spectra of AF complexes with different proteins in soluble (“Monomers”) and fibrillar (“Fibrils”) forms.

Thus, the dye can be used for the detection of amyloid-like beta-2 microglobulin, lysozyme, and insulin fibrils in solution. In addition, despite the fact that insulin is in soluble form (hexamer), its fluorescence peak is at 590 nm, which is not characteristic of the other AF/monomer complexes studied. However, the intensity of fluorescence in the complex with insulin fibrils grows by an order of magnitude compared to AF in complex with soluble insulin. This may be due to the presence of insulin in hexameric form.

Congo red is capable of forming supramolecular complexes and these complexes, at times, bind to amyloid-like fibrils [10,11]. The formation and stabilization of such complexes play a role in changes in dye spectral properties and the emergence of cooperative binding ability [12]. The notion of complex formation is supported by both computer simulation [13] and the binding stoichiometry of Congo-red-like dyes [5]. The change in the absorption spectra of such dyes can be associated with stabilization of molecule positions in complexes in the context of dye/beta-folded-structure interactions. The orientation of fluorophores, upon binding, can also lead to changes in spectral fluorescence characteristics, including an increase in quantum yield [14]. In this light, the data obtained regarding the insulin (soluble, hexameric)/AF complex is revealing. The lengths of beta pleated sheets are different between insulin hexamers and insulin fibrils, despite the ability of both to stabilize supramolecular complexes [15] [16] [17]. In addition, for insulin to form fibrils from hexamers, the protein must transition from the monomeric state to an oligomeric state [18]. Structures capable of stabilizing dye supramolecular complexes are present both in hexamers and in fibrils, and we postulate that the difference in fluorescence quantum yield is related to the difference in the lengths of dye supramolecular structures. Monomers of lysozyme and beta-2-microglobulin change the spectral properties of AF, but are not capable of stabilizing supramolecular complexes. Apparently, the pronounced changes in Congo-red-like dye spectral properties are connected with supramolecular complex length. Such lengths are directly dependent on the size of pleated sheet protein structures that promote supramolecular complex stability.

## Conclusion

The AF dye was synthesized, and it was shown that during interactions with beta-2 microglobulin, lysozyme, and insulin amyloid-like fibrils it undergoes changes in absorption spectrum and fluorescent properties. When amyloid-like fibrils are present, the intensity of the 590 nm fluorescence emission peak is increased for all of the proteins tested. The fluorescence emission intensity peak (FEIP) for monomeric beta-2-microglobulin and lysozyme (in AF complexes) were seen at 650 nm. The FEIP for insulin, however, was different; insulin’s FEIP was located at 590 nm, which is the same wavelength for all of the fibrillar form proteins tested, yet at a 10-fold lower intensity. Rational design of drugs capable of modulating protein oligomerization or preventing fibrillogenesis will undoubtedly require detailed data on dye structure and function, including intermolecular interactions. In addition to the practical importance of this research, study of Congo-red-analogues may be a prerequisite to determining the molecular mechanisms of amyloid-like fibril / dye interactions.

## Acknowledgments

We thank Dr. Dmitry V. Lebedev (NRC “Kurchatov Institute” - PNPI) for the fruitful discussion of the results.

